# Interpreting single-step genomic evaluations as mixed effects neural networks of three layers: pedigree, genotypes, and phenotypes

**DOI:** 10.1101/2022.07.18.500526

**Authors:** Tianjing Zhao, Hao Cheng

## Abstract

The single-step approach has become the most widely-used methodology for genomic evaluations when only a subset of phenotyped individuals in the pedigree are genotyped, where the genotypes for non-genotyped individuals are imputed based on gene contents of genotyped individuals through their pedigree relationships. We proposed a new method named single-step NN-MM to represent the single-step genomic evaluations as mixed effects neural networks of three sequential layers: pedigree, genotypes, and phenotypes, where the gene contents of non-genotyped individuals are sampled based on pedigree, genotypes, and phenotypes. In simulation analysis, the single-step NN-MM had similar or better prediction performance than the conventional single-step approach. In addition to imputation of genotypes using three sources of information including phenotypes, genotypes, and pedigree, single-step NN-MM provides a more flexible framework to allow nonlinear relationships between genotypes and phenotypes, and individuals being genotyped with different SNP panels. The single-step NN-MM has been implemented in a package called “JWAS”.

## Background

The single-step approach (Legarra *et al*. 2009; Christensen and Lund 2010; Fernando *et al*. 2014) has been successfully adopted in genomic evaluations when only a subset of phenotyped individuals in the pedigree are genotyped. The single-step approach utilizes information from genotyped and non-genotyped relatives in two equivalent ways: (a) calculating an improved relationship matrix from pedigree and observed genotypes to model the covariances of breeding values for all relatives (Legarra *et al*. 2009); or equivalently, (b) imputing genotypes for non-genotyped individuals linearly based on gene contents of genotyped individuals and the pedigree, and propagating the uncertainty from the imputation by fitting additional random effects accounting for imputation errors in genomic evaluations (Fernando *et al*. 2014) (see Appendix). In practice, the linear imputation in (b) can be obtained by modeling the gene content for each marker as a quantitative trait of very high heritability and fitting the “expected” gene content as random effects based on covariances defined by the pedigree (Gengler *et al*. 2007). Thus, from the latter interpretation (b) of the single-step approach, three sequential layers of information are involved: pedigree, genotypes, and phenotypes. This leads to our new representation of the single-step approach as a neural network, which consists of three fully-connected sequential layers of information: pedigree (input layer), genotypes (middle layer), and phenotypes (output layer), as demonstrated in Figure 1.

**Figure 1.**
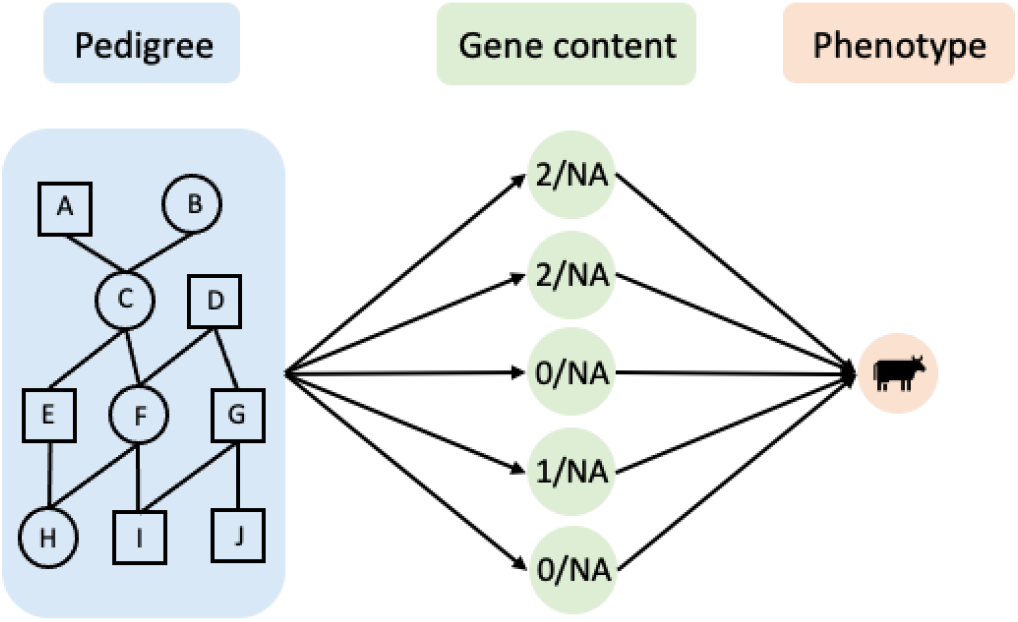
Framework of single-step NN-MM with three fully-connected sequential layers of data: pedigree, genotypes, and phenotypes. Between the layer of pedigree and the layer of genotypes, the gene content of each marker is treated as a quantitative trait, and the pedigree is used to define the random effects covariance matrix. Each node in the middle layer represents the gene content of one marker. “NA” denotes missing values. For example, the nodes in the middle layer may be 2,2,0,1,0 for a genotyped individual or all missing (“NA”) for a non-genotyped individual. For non-genotyped individuals, all gene contents are missing and will be sampled conditional on pedigree, genotypes, and phenotypes in MCMC.

We have proposed a method named “NN-MM”, the mixed effects neural network for quantitative genetics, to extend the mixed models (“MM”) to neural networks (“NN”) by adding intermediate layers of data (e.g., gene expression levels) between genotypes and phenotypes layers (Zhao *et al*. 2022, 2021). Better prediction accuracies were observed when intermediate omics data were incorporated into genomic prediction using NN-MM. In this paper, we will show that NN-MM can be adopted to incorporate pedigree, genotypes, and phenotypes information as a unified network named “single-step NN-MM”, thus providing a new representation of the single-step approach, and yielding equivalent or higher prediction accuracies, due to advantages described below.

There are some advantages of single-step NN-MM over the conventional single-step approach (Legarra *et al*. 2009; Christensen and Lund 2010; Fernando *et al*. 2014). Firstly, in the conventional single-step approach, gene contents of non-genotyped individuals are imputed based on the genotypes of genotyped individuals only through pedigree relationships. This can be considered as a pre-analysis processing using Gengler’s method (Gengler *et al*. 2007), and phenotypes are not included in this pre-analysis. We will show that, in single-step NN-MM, such pre-analysis is not needed, and gene contents of non-genotyped individuals are “imputed” based on pedigree, genotypes, and phenotypes in the Bayesian neural networks during Markov chain Monte Carlo (MCMC). Secondly, in single-step NN-MM, the relationships between genotypes and phenotypes can be approximated by nonlinear activation functions of the neural network to introduce non-linearity between genotypes and phenotypes. Lastly, the conventional single-step approach only allows individuals to be genotyped using the same SNP panel (i.e, same markers for all genotyped individuals), while single-step NN-MM can include individuals genotyped by different SNP panels (i.e., different markers for genotyped individuals) without pre-analysis.

In this paper, we will present single-step NN-MM for genomic evaluations, study its performance, and compare it to the conventional single-step approach (Legarra *et al*. 2009; Christensen and Lund 2010; Fernando *et al*. 2014). Here, we focus on studying the effect of fitting pedigree, genotypes, and phenotypes jointly as three unified fully-connected sequential layers, in which gene contents of non-genotyped individuals are sampled conditional on all three layers of data. Same assumptions of linearity and individuals being genotyped using the the same SNP panel, as in the conventional single-step approach, were used in singe-step NN-MM (i.e., a linear activation function, and individuals genotyped with the same SNP panel). We will also show that, in terms of predicting genetic values, single-step NN-MM is equivalent to the genomic BLUP (GBLUP) or the pedigree-based BLUP (PBLUP) when all or none of the phenotyped individuals in the pedigree are genotyped, respectively.

## Methods

In the single-step NN-MM, three sequential layers of information form a unified Bayesian neural network, as demonstrated in Figure 1. At each iteration of MCMC, unknowns will be sampled using Gibbs sampling from their full conditional posterior distributions as: (1) from the input layer (pedigree) to the middle layer (gene contents): PBLUP; (2) from the middle layer (gene contents) to the output layer (phenotypes): GBLUP or Bayesian Alphabet; and (3) sampling missing values in the middle layer (gene contents) based on three layers of information including pedigree, observed genotypes, and phenotypes.

### From input layer (pedigree) to middle layer (gene contents): Pedigree-based BLUP

Assuming there are *m* nodes in the middle layer (i.e., the number of markers is *m*), the gene content of *j*th SNP marker (*j* = 1,…, *m*) can be modeled as

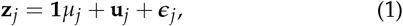

where **z**_*j*_ is a vector of (observed and sampled) gene contents of marker *j* for all individuals, *μ_j_* is its overall mean with a flat prior, **u**_*j*_ is the vector of gene content deviations with a prior 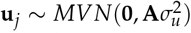, and **A** is the numerator relationship matrix calculated from pedigree. The random residual ***ϵ***_*j*_ is included to allow the use of mixed model equations, and to account for mistyping or pedigree errors (Gengler *et al*. 2007). The prior of ***ϵ***_*j*_ is 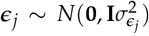. In principle, the heritability of the gene content should be 1 if the genotypes and pedigree information are perfectly correct, and a small value of estimated heritability indicates that there are problems in either genotypes or pedigree. Variance components are treated as unknowns in single-step NN-MM, and scaled inverse chi-square distributions are assigned as prior distributions for variance components.

### From middle layer (gene contents) to output layer (phenotypes): GBLUP or Bayesian Alphabet

The phenotypes can be modeled as:

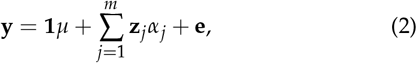

where **y** is the vector of phenotypes, *μ* is the overall mean with a flat prior, **z**_*j*_ is a vector of (observed and sampled) gene contents for the jth marker (*j* = 1,…, *m*), and *α_j_* is the corresponding marker effect. GBLUP (Habier *et al*. 2007; VanRaden 2008; Hayes *et al*. 2009) or Bayesian Alphabet (Meuwissen *et al*. 2001; Kizilkaya *et al*. 2010; Habier *et al*. 2011; Park and Casella 2008; Cheng *et al*. 2015; Gianola and Fernando 2019; Erbe *et al*. 2012; Moser *et al*. 2015; Cheng *et al*. 2018b) can be applied to sample marker effects or breeding values.

### Sampling missing values in the middle layer (gene contents)

For each marker, the missing genotypes will be sampled by Hamiltonian Monte Carlo (HMC) (Betancourt 2018) from its full conditional posterior distributions. Here we label the matrices related to non-genotyped and genotyped individuals with subscripts “n” and “g”, respectively. For *j*th marker, the full conditional posterior distribution of the missing gene content is proportional to the product of its prior and the likelihood:

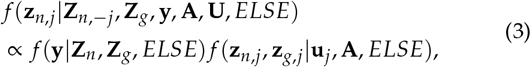

where ELSE denotes the current values of all other unknowns except **Z**_*n*_ = [**z**_*n*,1_,…, **z**_*n,m*_] and **U** = [**u**_1_,…, **u**_*m*_]. The **Z**_*n*,–*j*_ represents **Z**_*n*_ without jth column. All derivations are in the Appendix.

### Compared to the conventional single-step approach

Besides allowing non-linearity and individuals being genotyped with different SNP panels, the major difference between single-step NN-MM and the conventional single-step approach is on genotype imputation. In the conventional single-step approach, as described in Gengler *et al*. (2007), the gene content of each marker is treated as a quantitative trait as in equation (1), that is, the genotypes of non-genotyped individuals are imputed based on the observed genotypes of genotyped individuals through their pedigree relationships, and phenotypes are not included in the imputation (i.e., 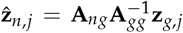 if *μ_j_* in equation (1) is known). This breaks down the three-layer unified network into two separate sub-networks: (1) genotype imputation: a sub-network from pedigree layer to genotypes layer, and (2) genomic prediction: a sub-network from genotypes layer to phenotypes layer. In single-step NN-MM, however, three layers of information are treated as a single unified network, such that the missing genotypes will be sampled based on three layers of information: pedigree, genotypes, and phenotypes, as shown in equation (3). Thus, the single-step NN-MM may have better prediction performance than the conventional single-step approach.

### Data Analysis

Given a linear relationship between genotypes and phenotypes, and the same SNP panel for all genotyped individuals are assumed in the conventional single-step approach, the same assumptions were used in single-step NN-MM (i.e., a linear activation function, and individuals genotyped with the same SNP panel) to compare the prediction performance of these two methods. In detail, GBLUP was used between the middle layer (gene contents) and the output layer (phenotypes) in single-step NN-MM (i.e., SS-NN-GBLUP), and its performance was compared to the conventional single-step GBLUP approach (i.e., SS-GBLUP).

A pig dataset from Cleveland *et al*. (2012) was used, which contains 3,534 genotyped individuals, and a pedigree of 6,473 individuals including parents and grandparents of the genotyped animals. The heritability of gene content for each marker is close to one (Forneris *et al*. 2015). For simplicity and computational efficiency, the first 100 SNPs were used. A random sample of five percent of these markers were selected as quantitative trait loci (QTL). Phenotypes were simulated with a heritability of 0.7 and a phenotypic variance of 1. The youngest 100 individuals, whose genotypes were observed but phenotypes were unknown, were used for testing, and the remaining individuals (i.e., 3434 individuals)with known phenotypes were used for training.

When all individuals were genotyped or non-genotyped, the performance of the single-step NN-MM were compared with the conventional GBLUP and PBLUP, respectively, using 10 replicates. Different proportions of non-genotyped individuals in the training dataset were considered, including 50%, 70%, 90%, and 99%. 10 replicates were performed for each scenario. The prediction accuracy was calculated as the Pearson correlation between the true breeding values and the estimated breeding values for individuals in the testing dataset. For both methods, 20,000 MCMC iterations were applied to ensure the convergence.

## Results

As shown in Figure 2, when all individuals were genotyped, the prediction accuracies of single-step NN-MM were almost identical to GBLUP. These results were expected as the single-step NN-MM was equivalent to a conventional linear mixed model in this scenario. When all individuals were non-genotyped, single-step NN-MM had almost identical prediction accuracy to PBLUP. These results were expected as the values of all nodes in the middle layer were missing, and only pedigree and phenotype information were used in single-step NN-MM.

**Figure 2.**
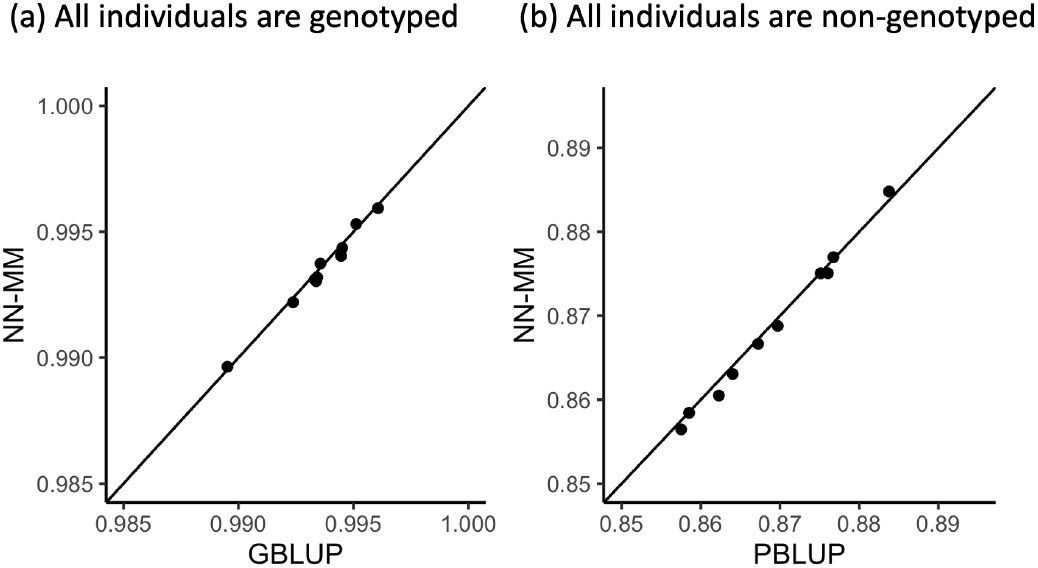
The prediction performance of single-step NN-MM with the linear activation function versus conventional GBLUP model, when all individuals are genotyped (a). The prediction performance of single-step NN-MM versus conventional PBLUP model, when all individuals are non-genotyped (b). 10 replicates were applied. The diagonal line was used for reference such that a dot close to this line represents a replicate with similar prediction accuracy of both methods.

The results when genotyping a proportion of phenotyped individuals were shown in Figure 3. Overall, the prediction accuracy of both methods decreased when the proportion of non-genotyped individuals increased. The SS-NN-GBLUP (red solid line) showed similar prediction accuracy as SS-GBLUP approach (blue solid line) in most scenarios, and a significant increase for SS-NN-GBLUP over SS-GBLUP approach was observed when 99% of individuals in the training datasets were not genotyped (pairwise t-test with P-value about 0.005).

**Figure 3.**
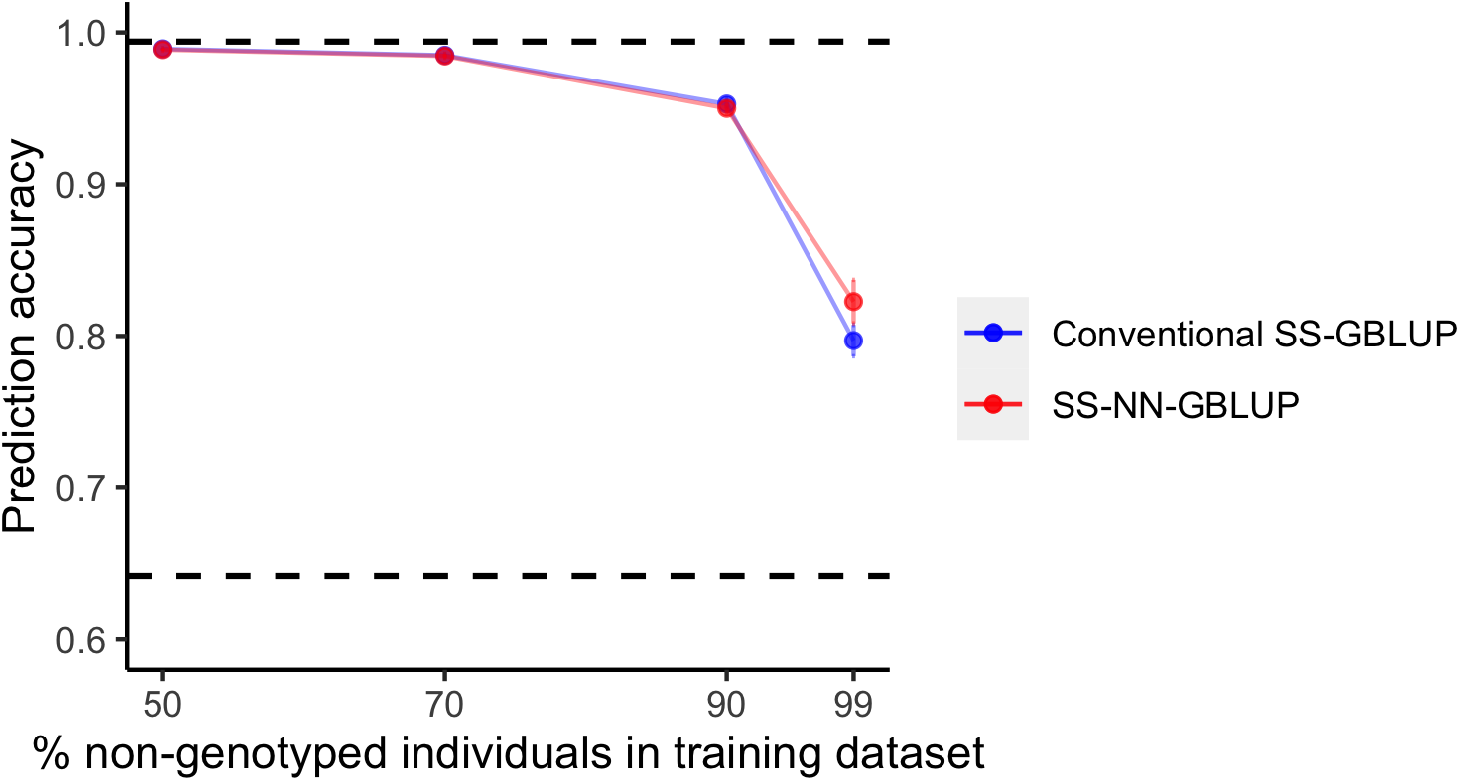
Prediction accuracies of single-step NN-MM with the linear activation function (SS-NN-GBLUP) in red solid line and the conventional single-step approach (SS-GBLUP) in blue solid line. Different proportions of non-genotyped individuals in the training dataset were considered, including 50%, 70%, 90%, 99%. The upper horizontal black dashed linear represents the conventional GBLUP model when all individuals were genotyped. The lower horizontal black dashed linear represents the conventional PBLUP model when no individuals were genotyped. Each dot represents the mean of prediction accuracies from 10 replicates, and the vertical bar is the mean ± its standard error.

## Discussion

In this paper, we proposed a new method named single-step NN-MM to represent the single-step genomic evaluations as mixed effects neural networks of three sequential layers: pedigree, genotypes, and phenotypes. Compared to the conventional single-step approach, the single-step NN-MM samples the gene contents of non-genotyped individuals based on phenotypes, pedigree and genotypes. Nonlinear relationships between genotypes and phenotypes are allowed, which can be approximated by nonlinear activation functions in the neural network. Also, single-step NN-MM can include individuals genotyped by different SNP panels (i.e., different markers for genotyped individuals). The single-step NN-MM has been implemented in a package called “JWAS” (Cheng *et al*. 2018a).

In our comparison, the same assumptions of linearity and identical SNP panels, as in conventional single-step approach, were used in singe-step NN-MM. Under these assumptions, when all individuals were genotyped or non-genotyped, single-step NN-MM had almost identical prediction performance as conventional linear mixed models (i.e., SS-NN-GBLUP versus GBLUP when all individuals were genotyped; SS-NN-MM versus PBLUP when all individuals were non-genotyped). Overall, when some individuals were not genotyped, single-step NN-MM had similar prediction accuracy as the conventional single-step approach, and the correlation between estimated marker effects from these two methods was high (see Table 1). A difference between singe-step NN-MM and the conventional single-step approach was observed when most individuals were not genotyped. Both conventional single-step approach and single-step NN-MM had significantly higher prediction accuracies compared to the GBLUP using genotyped individuals only (see Table 2).

**Table 1.** Correlation between marker effects estimated from conventional single-step approach and single-step NN-MM.

| % non-genotyped individuals |  |  |  |
| --- | --- | --- | --- |
| 50% | 70% | 90% | 99% |
| 0.999 (0.0003)<sup>a</sup> | 0.998 (0.001) | 0.993 (0.002) | 0.931 (0.015) |

<sup>a</sup> The average correlations from 10 replicates with the standard deviation in the bracket.

**Table 2.** The prediction performance comparison of excluding and including phenotypes of non-genotyped individuals.

| Method | % non-genotyped individuals |  |  |  |
| --- | --- | --- | --- | --- |
|  | 50% | 70% | 90% | 99% |
| GBLUP (genotyped individuals only) | 0.988<sup>a</sup> | 0.984 | 0.942 | 0.657 |
| conventional single-step | 0.989 | 0.985 | 0.953 | 0.797 |
| single-step NN-MM | 0.989 | 0.985 | 0.950 | 0.823 |

<sup>a</sup> The average prediction accuracies from 10 replications.

As we have described, in addition to allowing non-linearity and individuals being genotyped with different SNP panels, the major difference between single-step NN-MM and the conventional single-step approach is on genotype imputation. Besides genotypes and pedigree, phenotypic information is also used in the sampling of missing genotypes for non-genotyped individuals in single-step NN-MM. In most scenarios of our analysis, phenotypic information provided limited improvement for genotype imputation. We have performed our simulations with more SNPs (e.g., 200) and with SNPs randomly distributed across the genome, and similar patterns of results were observed. This may be due to at least two reasons. First, a single SNP only contributes to a small proportion of heritability, and the correlation between gene contents of one SNP and phenotypes is usually low. Secondly, phenotypes only help with the imputation of QTL (causal variants) and variants having high linkage disequilibrium with QTL. However, phenotypic information is used in the imputation of all SNPs including QTLs and markers. This may introduce errors to the imputation of markers. When genotypes of relatives provide limited information (e.g., most individuals are not genotyped), the additional benefits in genotype imputation by including phenotypic information is not negligible, and higher prediction accuracies were observed in single-step NN-MM.

In the current implementation of the single-step NN-MM, at each MCMC iteration, a naive multi-threaded parallelism (Bezanson *et al*. 2017) is used to employ multiple independent single-trait PBLUP models between input and middle layers in parallel. However, the speed improvement from multi-threaded parallelism is limited by the hardware (e.g., on a personal laptop). Ideally, with thousands of computer processors, the running time of multiple independent single-trait PBLUP models can equal the running time of one PBLUP model. Thus, more advanced parallel computing strategies needs to be further studied (e.g., Zhao *et al*. (2020); Breen *et al*. (2022)).

In summary, the single-step NN-MM method provides a novel and flexible framework for single-step genomic prediction. It allows “imputation” of genotypes for non-genotyped individuals using three sources of information: phenotypes, pedigree, and genotypes, in addition to nonlinear relationships between genotypes and phenotypes, and inclusion of individuals genotyped by different SNP panels.

## Availability of data and materials

Pig genotypes and pedigree used in the analysis are publicly available in Cleveland *et al*. (2012). The simulated phenotypes and all scripts are available at https://github.com/zhaotianjing/SSNNMM. The authors state that all data necessary for confirming the conclusions presented in the article are represented fully within the article.

## Author information

### Contributions

HC conceived the study. HC and TZ developed the methods, implemented the algorithms, planned the validations, wrote the manuscript. Both authors read and approved the final manuscript.

## Ethics declarations

### Ethics approval and consent to participate

Not applicable.

### Consent for publication

Not applicable.

## Appendix

### Conventional single-step approach as linear imputations

In conventional single-step approach, a pre-analysis processing is used to impute the gene contents of non-genotyped individuals from the genotypes of genotyped individuals through pedigree relationships. In detail, the *centered* gene content of each marker is treated as a normally-distributed quantitative trait as 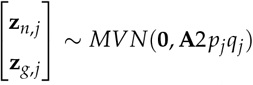, where *p_j_* is the allele frequency and *q_j_* = 1 – *p_j_*. Thus, the distribution of **z**_*n,j*_ conditional on **z**_*g,j*_ is a multivariate normal

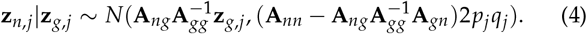

This can be written as

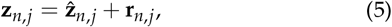

where 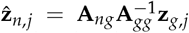 is the imputed genotypes for non-genotyped individuals, and **r**_*n,j*_ is the imputation uncertainty with 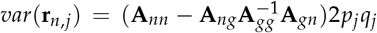. Extending above equation to multiple markers, we have 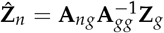.

After the pre-analysis processing, both imputed and observed genotypes will be used for genomic evaluation. An additional random effects will be used to account for the uncertainty from the genotype imputation. In detail, the phenotypes are written as

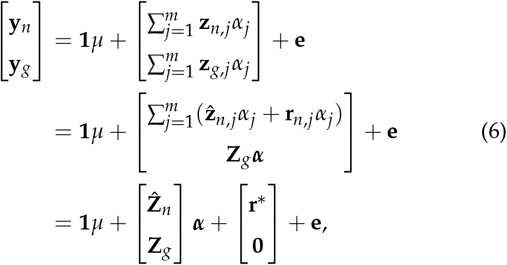

where *μ* is the overall mean, **Ẑ**_*n*_ is a matrix of imputed genotypes for non-genotyped individuals, **Z**_*g*_ is a matrix of observed genotypes for genotyped individuals, ***α*** is a vector of marker effects, and **e** is the random residuals with 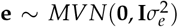. 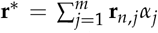 is a random vector to account for the sum of weighted imputation uncertainty with variance

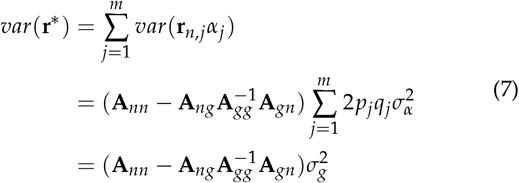

The conventional single-step GBLUP approach has been extended for Bayesian regression models to accommodate a wider assumptions of the marker effects (Fernando *et al*. 2014). It has been shown that the single-step Bayesian regression model with a normal prior for marker effects is equivalent to the single-step GBLUP approach in terms of predicting genetic values.

### Sample missing genotypes using Hamiltonian Monte Carlo

In single-step NN-MM, for *j*th marker (*j* = 1,…, *m*), the missing genotypes for non-genotyped individuals (**z**_*n,j*_) will be sampled by Hamiltonian Monte Carlo (HMC) (Betancourt 2018). In HMC, each unknown parameter **z**_*n,j*_ is paired with a “momentum” variable ***ϕ***_*n,j*_. The HMC constructs the Markov chain by a series of iterations. Following notations in Gelman *et al*. (2013), there are three steps in each iteration of the HMC:

1. updating the momentum variable independently of the current values of the paired parameter, i.e., ***ϕ***_*n,j*_ ~ *MVN*(**0**, **M**).
2. updating (**z**_*n,j*_, ***ϕ***_*n,j*_) via *L* leapfrog steps. In each leapfrog step, **z**_*n,j*_ and ***ϕ***_*n,j*_ are updated dependently and scaled by *t*. The leapfrog step below is repeated *L* times:

a. 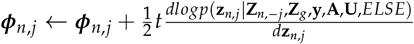;
b. **z**_*n,j*_ ← **z**_*n,j*_ + *t***M**^−1^***ϕ***_*n,j*_;
c. 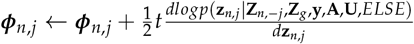.

The resulting state at the end of *L* repetitions will be denoted as 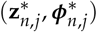.
3. calculating the acceptance rate, *r*, such that the resulting state will be accepted with probability *min*(1, *r*).

In our analyses, 10 leapfrog steps were applied in each HMC iteration (i.e., *L* = 10), the scale parameter *t* was 0.1, and **M** was set as an identity matrix.

Below we derive the gradient of the log full conditional posterior distribution of **z**_*n,j*_, which is required in HMC. For *j*th marker, the number of non-genotyped individuals denotes *n_n_*, and the full conditional posterior distribution of **z**_*n,j*_ is

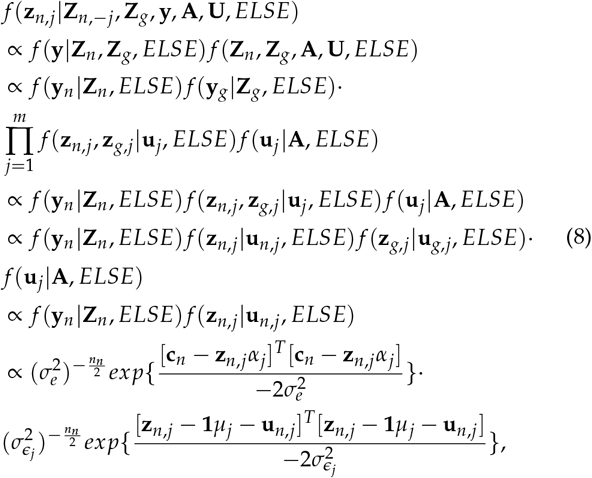

where **c**_*n*_ = **y**_*n*_ – **1***μ* – ∑_*j*′≠*j*_ **z**_*n,j*′_*α*_*j*′_, and *ELSE* denotes unknowns except **Z**_*n*_ = [**z**_*n*,1_,…,**z**_*n,m*_] and **U** = [**u**_1_,…,**u**_*m*_].

Thus, the log full conditional posterior distribution of **z**_*n,j*_ is

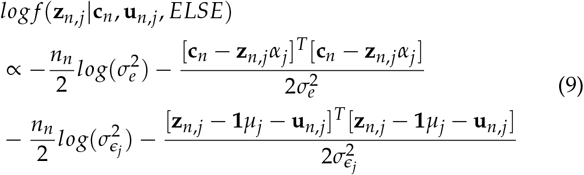

Thus, the gradient of the log full conditional posterior distribution of **z**_*n,j*_ is:

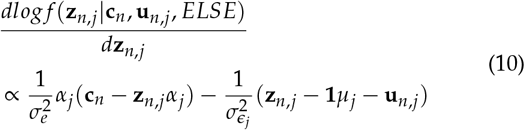

